# Identification of SARS-CoV-2 3CL Protease Inhibitors by a Quantitative High-throughput Screening

**DOI:** 10.1101/2020.07.17.207019

**Authors:** Wei Zhu, Miao Xu, Catherine Z. Chen, Hui Guo, Min Shen, Xin Hu, Paul Shinn, Carleen Klumpp-Thomas, Samuel G. Michael, Wei Zheng

## Abstract

The outbreak of coronavirus disease 2019 (COVID-19) caused by severe acute respiratory syndrome coronavirus 2 (SARS-CoV-2) has emphasized the urgency to develop effective therapeutics. Drug repurposing screening is regarded as one of the most practical and rapid approaches for the discovery of such therapeutics. The 3C like protease (3CL^pro^), or main protease (M^pro^) of SARS-CoV-2 is a valid drug target as it is a specific viral enzyme and plays an essential role in viral replication. We performed a quantitative high throughput screening (qHTS) of 10,755 compounds consisting of approved and investigational drugs, and bioactive compounds using a SARS-CoV-2 3CL^pro^ assay. Twenty-three small molecule inhibitors of SARS-CoV-2 3CL^pro^ have been identified with IC50s ranging from 0.26 to 28.85 μM. Walrycin B (IC_50_ = 0.26 µM), Hydroxocobalamin (IC_50_ = 3.29 µM), Suramin sodium (IC_50_ = 6.5 µM), Z-DEVD-FMK (IC_50_ = 6.81 µM), LLL-12 (IC_50_ = 9.84 µM), and Z-FA-FMK (IC_50_ = 11.39 µM) are the most potent 3CL^pro^ inhibitors. The activities of anti-SARS-CoV-2 viral infection was confirmed in 7 of 23 compounds using a SARS-CoV-2 cytopathic effect assay. The results demonstrated a set of SARS-CoV-2 3CL^pro^ inhibitors that may have potential for further clinical evaluation as part of drug combination therapies to treating COVID-19 patients, and as starting points for chemistry optimization for new drug development.

## Introduction

Coronavirus disease 2019 (COVID-19) has rapidly become a global pandemic since the first case was found in late 2019 in China. The causative virus was shortly confirmed as severe acute respiratory syndrome coronavirus 2 (SARS-CoV-2), which is a positive-sense single RNA virus consisting of four structural proteins and an RNA genome. Upon entering the host cell, the viral genome is translated to yield two overlapping polyproteins-pp1a and pp1ab ^1, 2^. 3CL protease (3CL^pro^, also known as main protease) of the related virus is excised from the polyproteins by its own proteolytic activity ^3^, and subsequently work together with papain-like protease to cleave the polyproteins to generate total 16 functional non-structural proteins (nsps). It was reported that the 3CL^pro^ of SARS specifically operates at 11 cleavage sites on the large polyprotein 1ab (790 kDa) ^3^, and no human protease has been found to share similar cleavage specificity ^4^. The cleaved nsps play essential roles in assembling the viral replication transcription complex (RTC) to initiate the viral replication. Although vaccine development is critically important for COVID-19, effective small molecule antiviral drugs are urgently needed. Because of its essential role and no human homolog, 3CL^pro^ is one of the most intriguing drug targets for antiviral drug development ^4, 5^. The inhibitors of 3CL^pro^ are most likely less-toxic to host cells ^4^.

Viral protease has been investigated as a drug target for decades resulting in several approved drugs for human immunodeficiency viruses (HIV) and hepatitis C virus (HCV) ^6^. Saquinavir was the first approved protease inhibitor for HIV, which started an era for this new class of antiviral drugs with the approval in 1995 ^6^. Saquinavir contains a hydroxyethylene bond, which functions as a peptidomimetic scaffold to block the catalytic function of the protease. Some of other approved protease inhibitors for HIV, such as ritonavir, nelfinavir, indinavir, lopinavir, amprenavir, atazanavir, fosamprenavir, and darunavir, share the similar inhibition mechanism with saquinavir ^6^. In addition to HIV, another group of approved viral protease inhibitors is used for treating HCV. Although the structure of HCV is different compared to HIV, the same function in cleaving viral precursor proteins renders the HCV protease a valid target for antiviral drug development. Asunaprevir, boceprevir, simeprevir, paritaprevir, vaniprevir, telaprevir, and grazoprevir, etc. have been approved by FDA for treatment of HCV ^6^. Regarding the coronavirus, the effort to target the SARS-CoV and SARS-CoV-2 3CL^pro^ has identified several drug candidates ^7^. However, to date, no 3CL^pro^ inhibitor has been specifically approved for SARS-CoV or SARS-CoV-2.

In this study, we have employed a SARS-CoV-2 3CL^pro^ assay that uses a self-quenched fluorogenic peptide substrate for a quantitative high throughput screen (qHTS) ^8^ of 10,755 compounds including approved drugs, clinically investigated drug candidates, and bioactive compounds. We report here the identification of 23 3CL^pro^ inhibitors with the IC_50_ ranging from 0.3 to 30 μM. The results from this study can contribute to the design of the synergistic drug combinations for treatment of COVID-19 as well as new starting points for lead optimization of SARS-CoV-2 3CL^pro^ inhibitors.

## Results

### Optimization of enzyme and substrate concentrations for 3CL^pro^ enzyme assay

To carry out a qHTS of SARS-CoV-2 3CL^pro^ inhibitors, a fluorogenic protease enzyme assay was used and optimized. As illustrated in **Figure 1**, the C-terminal of a peptide substrate links to a fluorophore (Edans) and the N-terminal has a fluorescence quencher (Dabcyl) that quenches the fluorescence signal of Edans. When the 3CL^pro^ hydrolyzes the substrate to yield two fragments, Dabcyl is separated with Edans, which relieves the fluorescence quenching effect resulting in an increase of fluorescence signal.

**Figure 1.**
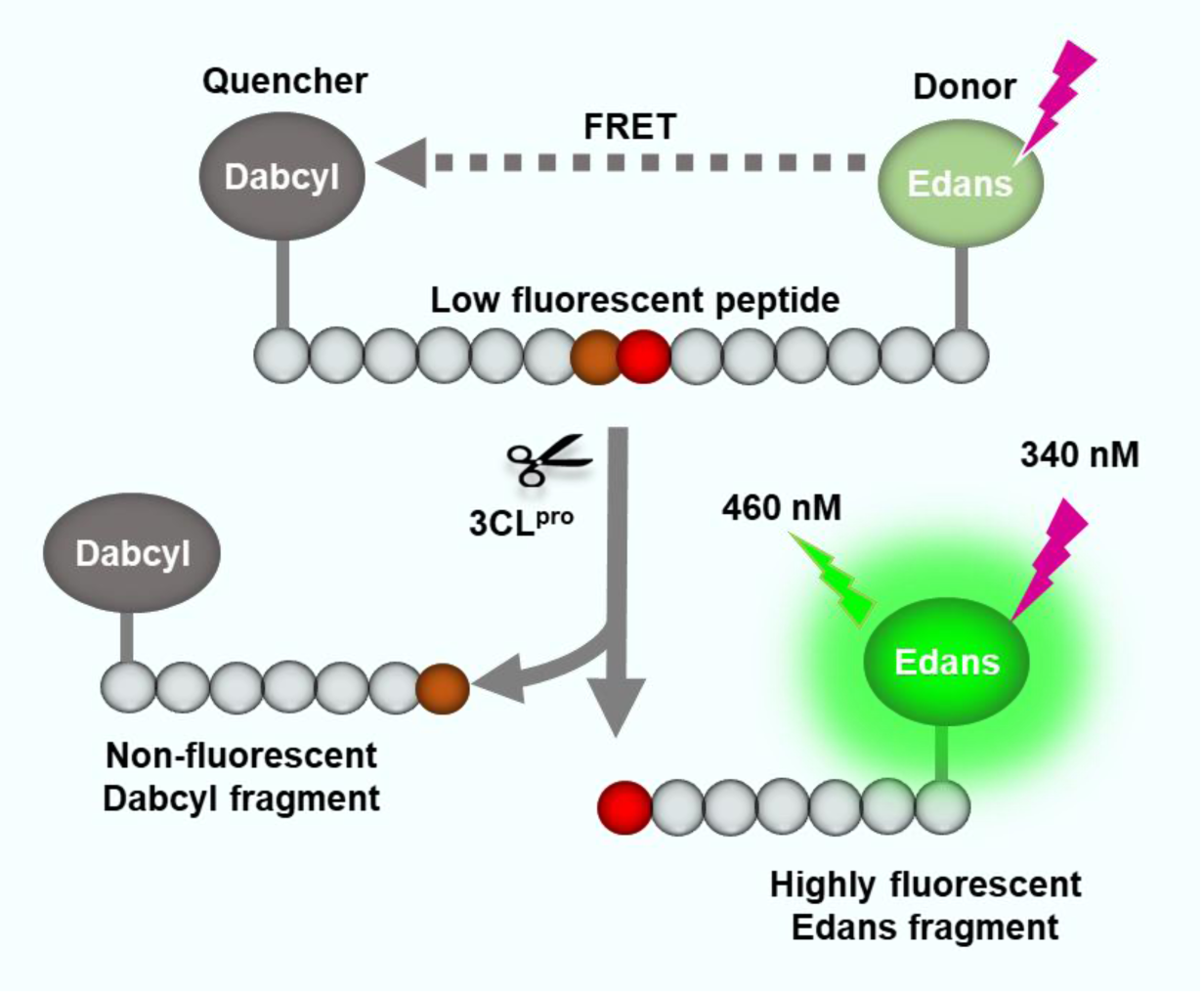
Schematic representation of the fluorogenic SARS-CoV-2 protease enzymatic assay. The peptide substrate exhibits low fluorescent because the fluorescence intensity of Edans in the C-terminal is quenched by the Dabcyl in the N-terminal of the substrate. The protease cleaves the substrate which breaks the proximity of the quencher molecule Dabcyl with the fluorophore Edans, resulting in an increase in fluorescence signal. This increase in fluorescence signal is proportional to the protease activity.

To optimize the assay conditions, different enzyme concentrations were first examined in this enzyme assay at 20 µM substrate concentration in a 384-well plate (**Figure 2a**). The signal-to-basal (S/B) ratio increased with the incubation times for all 3 enzyme concentrations tested (**Figure 2b**). The 120 min incubation resulted in S/B ratios of 2.4, 3.8, and 6.0-fold for enzyme concentrations of 25, 50 and 100 nM, respectively. We selected 50 nM enzyme concentration as an optimized condition for subsequent experiments.

**Figure 2.**
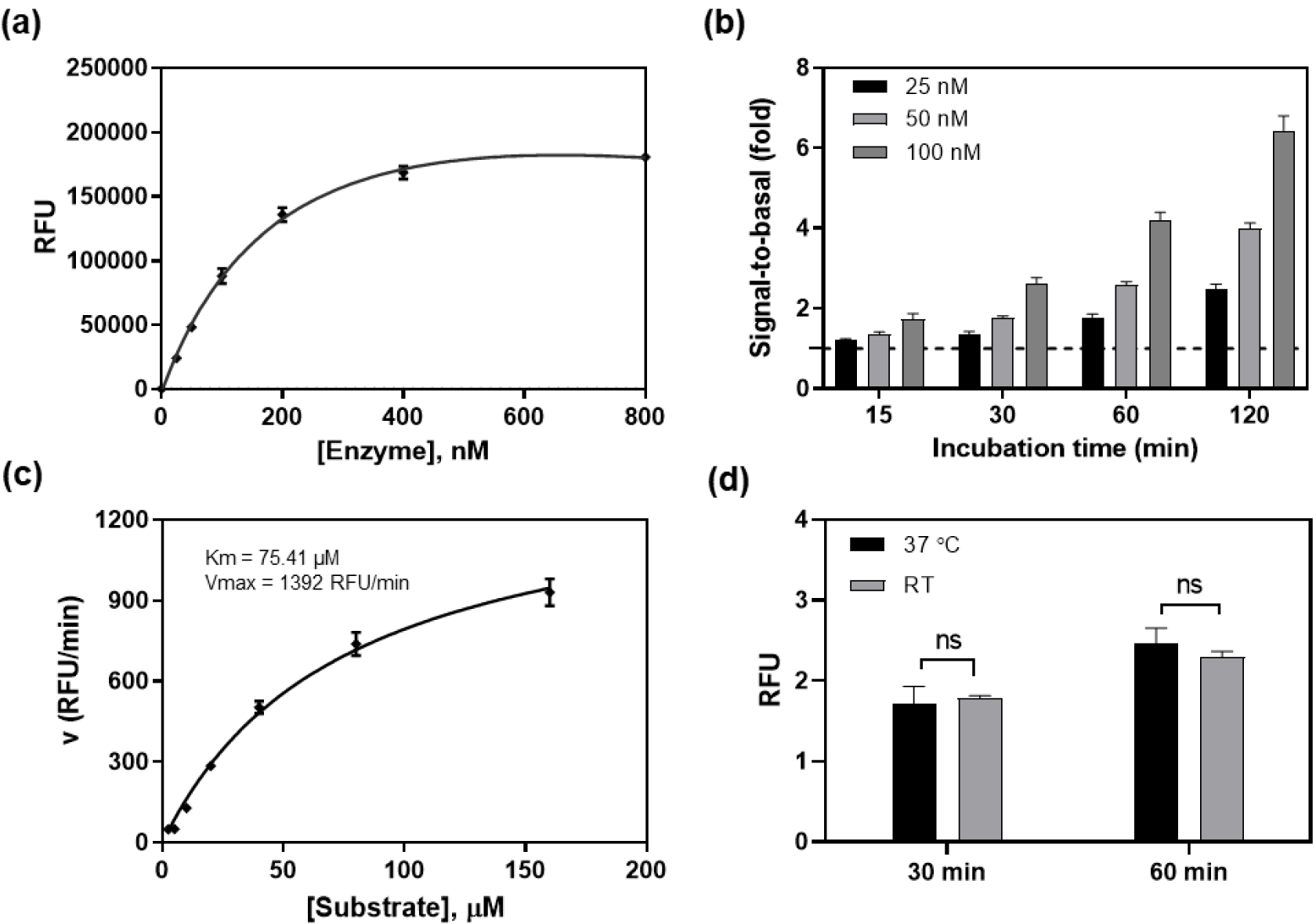
SARS-CoV-2 3CL^pro^ enzyme assay optimization. (a) Concentration-response curve of enzyme titration. With a fixed concertation of substrate (20 μM), the fluorescent intensity increased with enzyme concentrations. The linear response was observed at low enzyme concentrations. Measurement was conducted 2 h after initiating the reaction at RT. (b) The signal-to-basal (S/B) ratios of three enzyme concentrations within the linear range, at various incubation times. Dotted line represents the S/B = 1. (c) Enzyme kinetics. Michealis-Menton plot exhibited a Km of 75.41 μM and Vmax of 1392 RFU/min for SARS-CoV-2 3CL^pro^. (d) The S/B ratios determined at RT and 37 °C. No difference was observed in 1 h incubation between the two temperatures.

Enzyme kinetic study was then conducted to determine the Km and Vmax of this viral protease. Substrate concentrations ranging from 2.5 to 160 µM were used in this experiment with a fixed 50 nM enzyme (**Figure 2c**). The Km was 75.41 µM and the Vmax was 1392 RFU/min for this recombinant SARS-CoV-2 3CL^pro^. For a consideration of assay sensitivity, it is desirable to use the lowest enzyme concentration and substrate concentration (ideally under Km value) that still yield reliable S/B ratio (assay window, usually > 2-fold). Because inhibitors usually compete with substrates for binding to the free enzyme, a high substrate concentration can reduce potencies (IC50s) of inhibitors determined in enzyme assays ^9^. Thus, 50 nM 3CL^pro^ and 20 µM substrate were selected as the optimized conditions for qHTS. We also found that the assay performance at 37 °C was similar to that at RT. Therefore, the subsequent compound screens were conducted at RT (**Figure 2d**).

### Validation of 3CL^pro^ enzyme assay with a known protease inhibitor and measurements of HTS assay parameters

To validate the enzyme assay, a known SARS-CoV-2 3CL^pro^ inhibitor, GC376 ^10^, was evaluated in 1536-well plate format. GC376 concentration-dependently inhibited the enzyme activity of SARS-CoV-2 3CL^pro^ with an IC_50_ value of 0.17 µM. The highest tested concentration of GC376 (57.5 μM) exhibited a complete inhibition, where the fluorescent intensity was equal to the background substrate in the absence of enzyme. This IC_50_ value of GC376 is comparable to the reported value ^10^, indicating the reliability of this enzymatic assay (**Figure 3a**).

**Figure 3.**
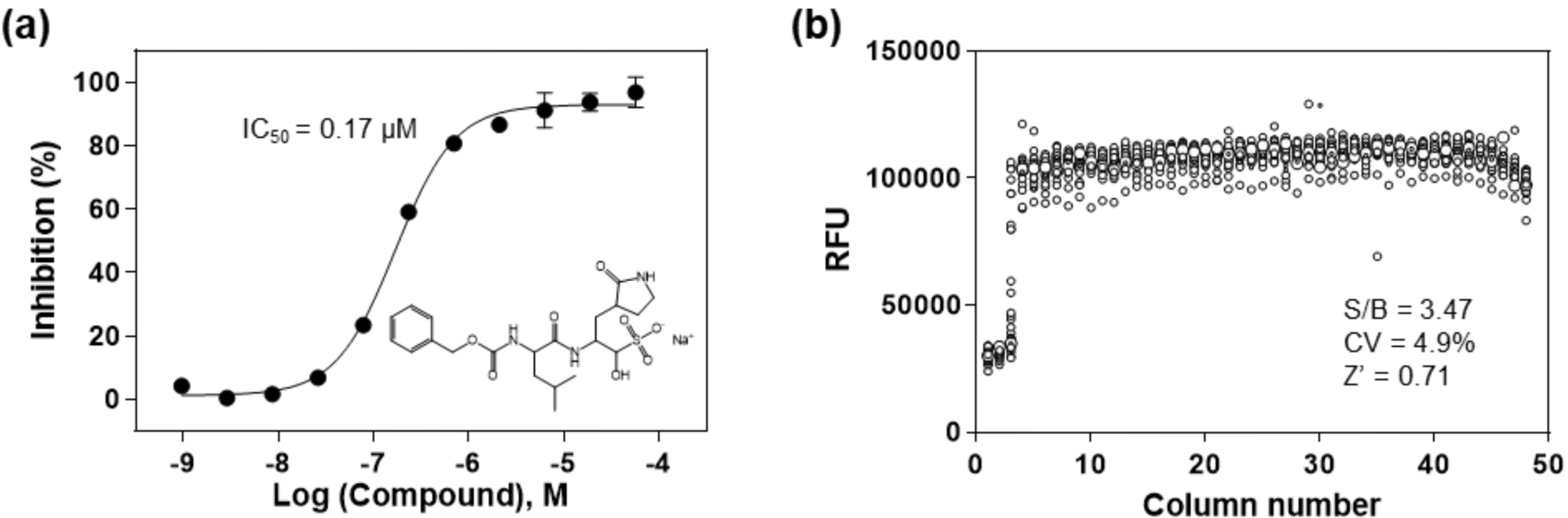
(a) Concentration response of the known 3CL^pro^ inhibitor, GC376. An IC_50_ of 0. 17 µM was determined for the inhibition of SARS-CoV-2 3CL^pro^. The substrate concentration was 20 μM and enzyme concentration was 50 nM in this experiment. (b) Scatter plot of the results from a DMSO plate in the 3CL^pro^ enzymatic assay in a 1536-well plate, where columns 1 and 2 in the plate contain substrate only, column 3 includes GC376 titration (1:3 dilution series from 57.5 µM), and columns 5-48 contain DMSO (23 nL DMSO in 4 µL reaction solution).

Since DMSO is the solvent for all compounds in our compound libraries, we tested a DMSO plate in 1536-well plate for the assessment of HTS assay parameters. A S/B ratio of 3.47-fold, a coefficient of variation (CV) of 4.9% and a Z’ factor of 0.71 were obtained in the 1536-well DMSO plate test, indicating this enzyme assay is robust for HTS (**Figure 3b**).

### Drug repurposing screen for 3CL^pro^ enzyme inhibitors

A primary screen of 10,755 compounds in the libraries containing approved drugs, investigational drug candidates, and bioactive compounds yielded 161 hits in which the hit rate was 1.5 % (Figure S1). Since the primary screen was done in four compound concentrations, compounds in dose-response curve classes 1-3 were selected as hits from the primary screen ^8, 11^. These primary hits were then “cherry-picked” for confirmation test in the same enzyme assay. Hit confirmation was performed at 11 compound concentrations at 1:3 titration. The confirmed hits were selected using a cutoff of maximal inhibition greater than 60 % and IC_50_ less than 30 µM. The results of primary screening and hit confirmation have been uploaded into the NCATS Open Data Portal for public access ^12^.

Because this SARS-CoV-2 3CL^pro^ assay is a fluorogenic assay, compounds with fluorescence quenching properties can suppress the fluorescence signal generated by the protease activity. To eliminate the false positives, we conducted a counter screen to identify compounds that quench the fluorescence of SGFRKME-Edans, the product of the 3CL^pro^ enzyme reaction, in the absence of the protease enzyme. Based on the standard curve (Figure S2), the 3CL^pro^ assay conditions generated signals that matched 2.085 μM of Edans fragment. Results indicated that 23 compound showed relatively negligible fluorescence quenching effect (**Table 1**). The concentration response curves of 6 most potent inhibitors of SARS-CoV-2 3CL^pro^ are shown in **Figure 4**.

**Table 1.** Activity of 20 identified compounds against SARS-CoV-2 3CL^pro^, and their CPE and cytotoxicity.

| Compound Name | 3CL<sup>pro</sup> Inhibition |  | SARS-CoV-2 CPE |  | Vero E6 Cytotoxicity |  |
| --- | --- | --- | --- | --- | --- | --- |
|  | IC<sub>50</sub> (μM) | Max Resp (%) | EC<sub>50</sub> (μM) | Efficacy (%) | CC<sub>50</sub> (μM) | Efficacy (%) |
| Walrycin B | 0.26 | 86.6 | 3.55 | 51.43 | 4.25 | 99.67 |
| Hydroxocobalamin | 3.29 | 89.56 | N/A, >20 | 0 | N/A, >20 | 0 |
| Suramin sodium | 6.5 | 99.49 | N/A, >20 | 0 | N/A, >20 | 0 |
| Z-DEVD-FMK | 6.81 | 90.48 | N/A, >20 | 0 | N/A, >20 | 0 |
| LLL-12 | 9.84 | 82.98 | N/A, >20 | 0 | 1.77 | 100 |
| Z-FA-FMK | 11.39 | 88.74 | 0.13 | 104.84 | N/A, >20 | 0 |
| DA-3003-1 | 2.63 | 70.64 | 4.47 | 53.77 | 7.74 | 112 |
| CAY-10581 | 9.2 | 60.62 | N/A, >20 | 0 | N/A, >20 | 0 |
| Fascaplysin | 9.96 | 63.44 | N/A, >20 | 0 | 1.26 | 99.28 |
| MG-115 | 12.7 | 74.89 | 0.023 | 72.005 | 1.13 | 115.48 |
| beta-Lapachone | 13.33 | 60.88 | N/A, >20 | 0 | 12.59 | 15.06 |
| Sepantronium bromide | 13.6 | 61.79 | N/A, >20 | 0 | 7.94 | 20.12 |
| NSC 95397 | 17.93 | 98.7 | N/A, >20 | 0 | 14.14 | 111.33 |
| Vitamin B12 | 18.02 | 71.06 | N/A, >20 | 0 | N/A, >20 | 0 |
| 4E1RCat | 18.28 | 76.33 | N/A, >20 | 0 | N/A, >20 | 0 |
| TBB | 20.49 | 77.96 | 14.13 | 28.29 | 8.91 | 80.93 |
| GW-0742 | 22.27 | 126.23 | N/A, >20 | 0 | N/A, >20 | 0 |
| Agaric acid | 23.54 | 110.16 | N/A, >20 | 0 | N/A, >20 | 0 |
| Oritavancin (diphosphate) | 24.15 | 93.29 | N/A, >20 | 0 | 14.13 | 23.51 |
| MK 0893 | 24.33 | 95.77 | 3.16 | 23.89 | 12.59 | 80.04 |
| SU 16f | 24.97 | 87.95 | N/A, >20 | 0 | 7.94 | 23.57 |
| Penta-O-galloyl-β-D-glucose hydrate | 27.77 | 109.94 | 12.59 | 28.86 | 11.22 | 80.28 |
| SP 100030 | 28.85 | 64.84 | N/A, >20 | 0 | N/A, >20 | 0 |

**Figure 4.**
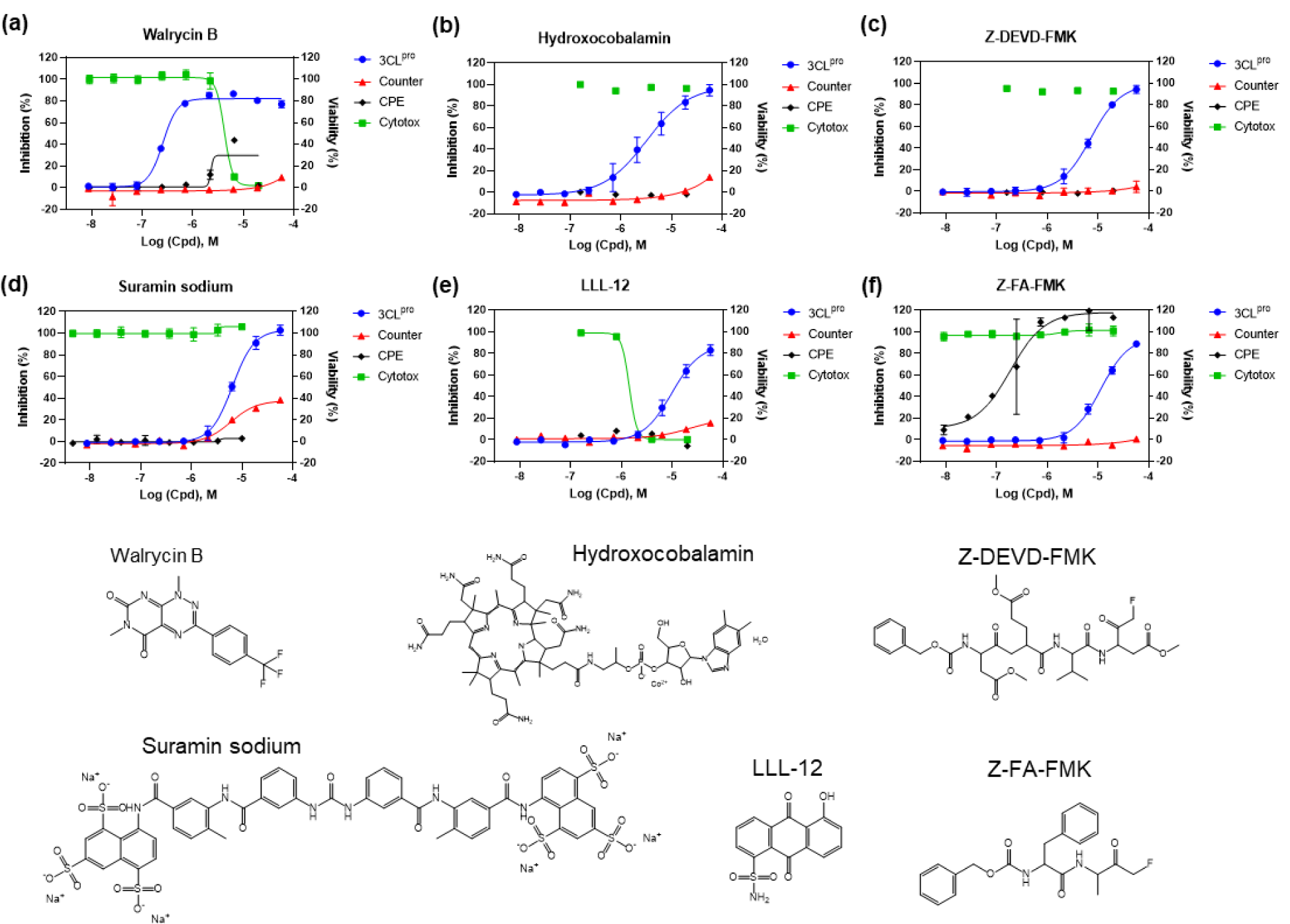
Concentration-response curves of the 6 most potent compounds with IC_50_ values < 15 μM and maximal inhibition > 80% determined in the SARS-CoV-2 3CL^pro^ enzyme assay. Enzyme assay (blue) and counter screen (red) curves correspond to left y-axis showing inhibitory results, CPE (black) and cytotoxicity (green) curves correspond to right y-axis showing cell viability. (a) Walrycin B, IC_50_ = 0.26 μM. (b) Hydroxocobalamin, IC_50_ = 3.29 μM. (c) Z-DEVD-FMK, IC_50_ = 6.81 μM. (d) Suramin sodium, IC_50_ = 6.5 μM. (e) LLL-12, IC_50_ = 9.84 μM. (f) Z-FA-FMK, IC_50_ = 11.39 μM. Primary CPE and cytotoxicity screens were conducted in 4 concentrations, only the hits were further confirmed with 8 concentrations with dilution ratio of 1:3.

### Confirmation of antiviral activity of 3CL^pro^ inhibitors in a SARS-CoV-2 live virus assay

To evaluate the antiviral effect of these 3CL^pro^ inhibitors against infections of SARS-CoV-2 virus, we tested the confirmed inhibitors in a cytopathic effect (CPE assay). Amongst the 6 most potent compounds in the 3CL^pro^ enzyme assay (**Figure 4**), walrycin B and Z-FA-FMK showed the rescue of SARS-CoV-2 induced CPE with the efficacies of 51.43% and 104.84%, respectively. The protease inhibitor Z-FA-FMK inhibited viral CPE with an EC_50_ of 0.13 μM, with no apparent cytotoxicity. Hydroxocobalamin, suramin sodium, and Z-DEVP-FMK were neither effective in CPE assay, nor cytotoxic to Vero E6 cells. Walrycin B and LLL-2 showed apparent toxicity with CC50 values of 4.25 µM and 1.77 μM, and full cytotoxicity levels. For other compounds identified, DA-3003-1, MG-115, TBB, MK0983, and Penta-O-galloyl-β-D-glucose hydrate exhibited CPE activity as well (**Table 1 and Figure S3**).

In addition, 9 compounds with partial quenching effect rescued cells from SARS-CoV-2 (**Figure 5** and **Table 2**). In these compounds, anacardic acid and AMG-837 showed the best rescue effect with the efficacy of 89.62% and 106.31%, respectively. Meanwhile, all of these compounds showed more or less cytotoxicity that renders an uncertainty of their anti-SARS-CoV-2 activity in cells. The response curves for other compounds with quenching effects can be found in **Figure S4**.

**Table 2.** Activity of 9 CPE active quenching compounds against SARS-CoV-2 3CL^pro^, and their CPE and cytotoxicity.

| Compound Name | 3CL<sup>pro</sup> Inhibition |  | SARS-CoV-2 CPE |  | Vero E6 Cytotoxicity |  |
| --- | --- | --- | --- | --- | --- | --- |
|  | IC<sub>50</sub> (μM) | Max Resp (%) | EC<sub>50</sub> (μM) | Efficacy (%) | CC<sub>50</sub> (μM) | Efficacy (%) |
| Anacardic Acid | 11.04 | 105.9 | 3.98 | 89.62 | 14.13 | 72.92 |
| Adomeglivant | 19.18 | 107.72 | 3.16 | 49.2 | 14.13 | 36.97 |
| Eltrombopag<br>olamine | 21.52 | 92.58 | 8.91 | 74.39 | 7.94 | 68.67 |
| GSK-3965 | 21.52 | 108.24 | 3.55 | 22.51 | 14.13 | 82.34 |
| GW5074 | 21.52 | 63.16 | 11.22 | 63.3 | 11.22 | 20.1 |
| Hexachlorophene | 21.52 | 117.06 | 0.79 | 43.57 | 4.47 | 63.02 |
| MK-886 | 21.52 | 109.11 | 11.22 | 69.18 | 14.13 | 40.83 |
| AMG-837 | 24.15 | 65.79 | 11.22 | 106.31 | 15.85 | 20.2 |
| MG-149 | 27.1 | 96.26 | 12.59 | 70 | 14.13 | 23.42 |

**Figure 5.**
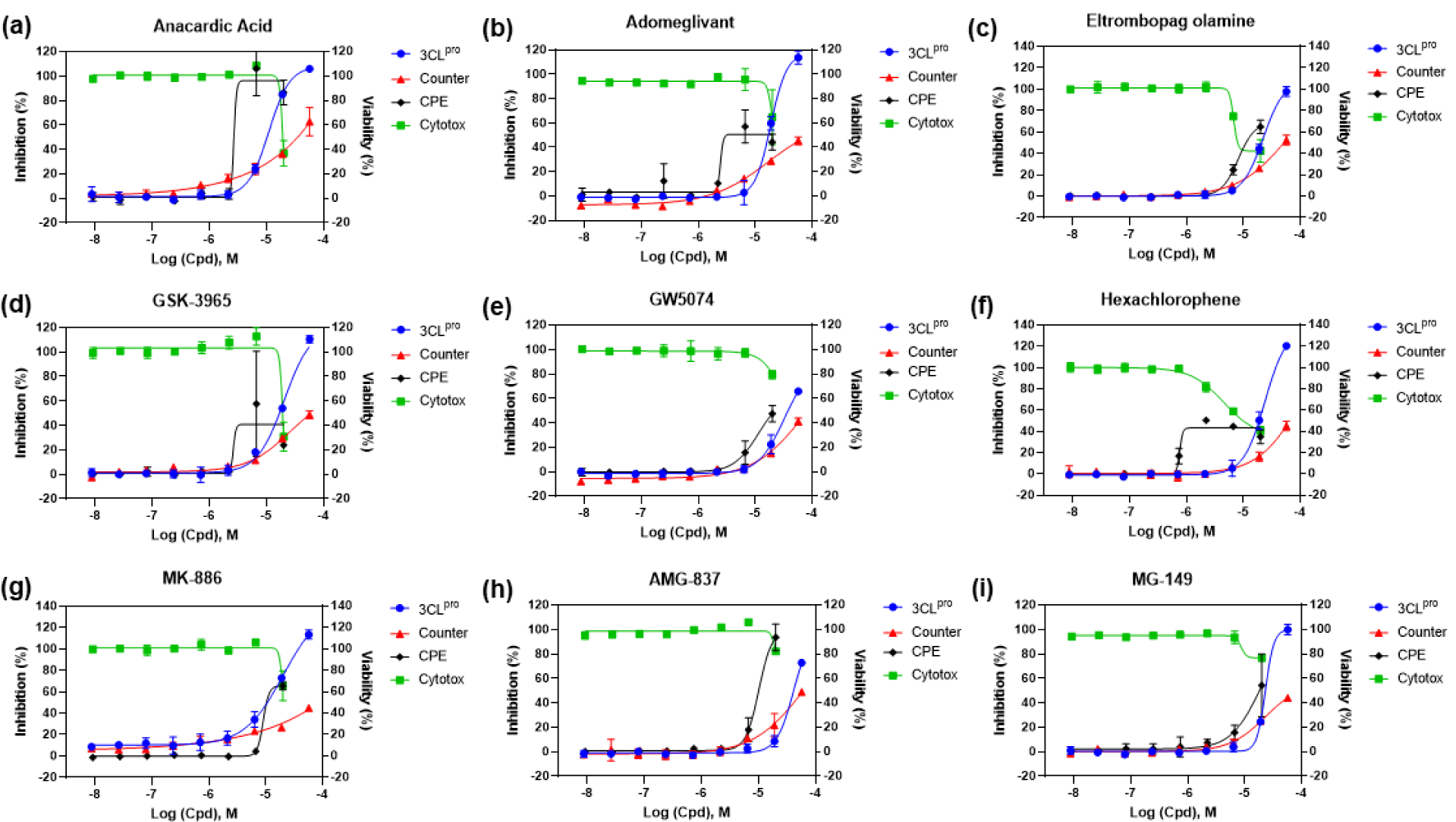
Concentration-response curves of the 9 compounds with partial quenching effect and CPE activity. (a) Anacardic Acid, CPE efficacy = 89.62%. (b) Adomeglivant, CPE efficacy = 49.2%. (c) Eltrombopag olamine, efficacy = 74.39%. (d) GSK-3965, efficacy = 22.51. (e) GW574, CPE efficacy = 63.3%. (f) Hexachlorophene, CPE efficacy = 47.57%. (g) MK-886, CPE efficacy = 69.18%. (h) AMG-837, CPE efficacy = 106.31%. (i) MG-149, CPE efficacy = 70%.

Interestingly, there was poor correlation between the 3CL^pro^ enzyme and CPE assays. This could be due to cytotoxicity of compounds obscuring CPE effects in live cells. The reason for weak potencies of these compounds in the CPE assay than in the 3CL^pro^ enzyme assay could be caused by unoptimized drug properties such as poor cell membrane permeability of compounds. Conversely, some compounds, such as Z-FA-FMK, exhibited more potent activities in the CPE assay than these in the 3CL^pro^ enzyme assay that may be due to polypharmacology (i.e. targeting multiple steps in viral replication process).

### Modeling analysis

We docked the identified compounds to the active site of 3CL^pro^ to further investigate their potential binding to the protease target. The peptide-like inhibitors Z-DEVD-FMK and Z-FA-FMK fit well in the active site of 3CL^pro^ by forming a covalent bond between the keto and the catalytic residue Cys145, while the side groups were orientated to the S1, S2 and S4 pocket (**Figure 6**). Small molecule inhibitors like Walrycin B and LLL-12 were found to bind to the S1 or S1’ pocket near Cys145, but no specific binding interactions were observed. Most of other compounds such as Suramin sodium and Hydroxocobalamin did not dock in the active site of 3CL^pro^ and appeared not to be protease inhibitors.

**Figure 6.**
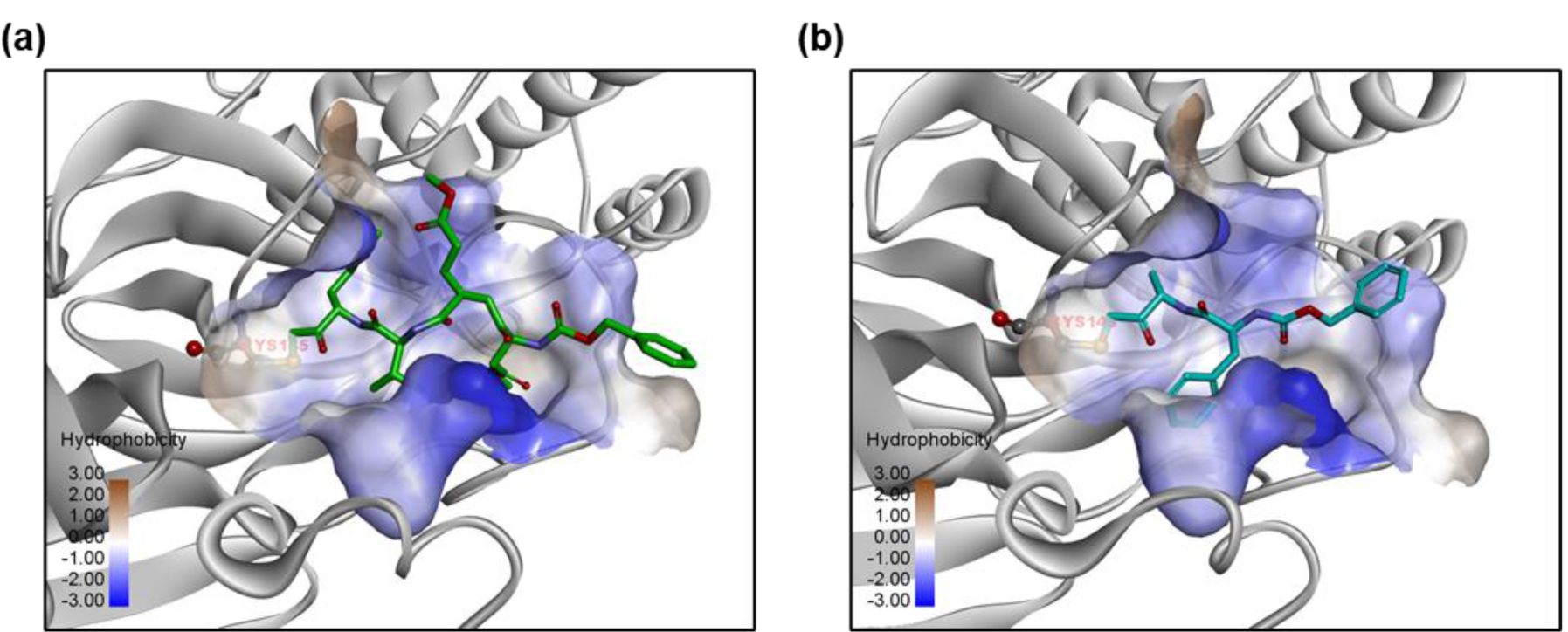
Predicted binding models of (a) Z-DEVD-FMK and (b) Z-FA-FMK bound to the active site of 3CL^pro^. The protein 3CL^pro^ (grey) is represented in ribbons and the active site is shown with the hydrophobic protein surface. Small molecule inhibitors are shown in sticks. The catalytic residue Cys145 in the binding pocket is highlighted.

## Discussion

Viral protease is a valid antiviral drug target for RNA viruses including coronaviruses ^13^. In response to the COVID-19 pandemic, great efforts have been made to evaluate the possibility of repurposing approved viral protease inhibitor drugs for the clinical treatment of the disease. Unfortunately, the combination of lopinavir and ritonavir, both approved HIV protease inhibitors, failed in a clinical trial without showing benefit compared to the standard of care ^14^. To address this unmet need, several virtual screens and a drug repurposing screen were performed to identify SARS-CoV-2 3CL^pro^ inhibitors. Ma et al. screened a focused collections of protease inhibitors using an enzyme assay and identified Boceprevir, GC376, and three calpain/cathepsin inhibitors as potent SARS-CoV-2 3CL^pro^ inhibitors ^10^. Amongst them, Boceprevir, an FDA-approved HCV drug, not only showed the inhibition of 3CL^pro^ with an IC_50_ of 4.13 μM, but also has an EC_50_ of 1.90 μM against SARS-CoV-2 virus infection in the CPE assay ^10^. Lopinavir and ritonavir did not show inhibition to 3CL^pro^ in the study that indicated both of them have weak inhibitory activity against SARS-CoV-2 3CL^pro^.

In the virtual screen efforts, novel compounds were designed and synthesized by analyzing the substrate-binding pocket of SARS-CoV-2 3CL^pro^. The two lead compounds, 11a and 11b, designed by Dai et al. presented high potency in both enzyme inhibition and anti-SARS-CoV-2 infection activity ^15^. In another more comprehensive study, Jin et al. identified six compounds that have IC_50_ values ranging from 0.67 to 21.4 µM against SARS-CoV-2 3CL^pro^ by applying structure-assisted drug design and compound library repurposing screen ^2^.

In our qHTS of 10,755 compounds using the 3CL^pro^ enzyme assay, we identified 27 compounds that inhibited SARS-CoV-2 3CL^pro^ enzymatic activity. Among the most potent compounds(**Figure 4**), walrycin B (IC_50_ = 0.27 μM) is the most potent inhibitor found in this screen. Walrycin B is an analog of toxoflavin (a phytotoxin from *Burkholderia glumae*) with potent activity of inhibiting bacteria growth. It was named as Walrycin B because it was found to inhibit the WalR activity in bacteria ^16^. The WalK/WalR two-component signal transduction system is essential for bacteria cell viability.

Hydroxocobalamin is a synthetic vitamin B12 (cobalamin) that is used in the clinics via intravenous administration. The antiviral effect of Vitamin B12 on HIV and HCV was reported previously. It was reported that Vitamin B12 inhibited the HIV integrase ^17^. Li et al. reported that Vitamin B12 inhibited the HCV protein translation via the inhibition of HCV internal ribosome entry site ^18^. We found that hydroxocobalamin inhibited the SARS-CoV-2 3CL^pro^ activity in this study.

Suramin is an FDA-approved anti-parasitic drug for trypanosomiasis and onchocerciasis and has to be given by intravenous injection as it has poor bioavailability when being taken orally. The IC_50_ for inhibition of SARS-CoV-2 3CL^pro^ is 6.5 μM, which is much lower than the reported human plasma concentration of 97 - 181 μM (126 – 235 μg/ml) ^19^. Suramin is an old drug with extensive polypharmacology ^20^. Broad antiviral effects of suramin were reported including HIV, Dengue virus, Zika virus, Ebola virus, Hepatitis B and C viruses, Herpes simplex virus, Chikungunya virus, and Enterovirus ^20^. The antiviral activity of suramin may be exerted through inhibiting viral entry and replication. Suramin can efficiently inhibit Chikungunya virus and Ebola envelope-mediated gene transfer to host cells ^21^. Multiple studies have also revealed that suramin interferes with viral RNA synthesis by targeting viral RNA-dependent RNA polymerase ^22, 23^. A recent study by de Silva et al. proposed suramin might prevent SARS-CoV-2 viral entry into cells^24^. Different from these previously reported various mechanisms of action, we found that suramin also targeted 3CL^pro^ enzyme of SARS-CoV-2, which is a new mechanism of action for this drug.

Another 3CL^pro^ inhibitor identified is Z-DEVD-FMK (IC_50_ = 6.81 μM). It is a cell permeable fluoromethyl ketone (FMK)-derivatized peptide acting as an irreversible caspase 3 inhibitor. It has been extensively studied as a neuroprotective agent as it inhibited caspase 3 induced apoptotic cell death in acute neurodegeneration ^25, 26^. Another similar peptide-like inhibitor Z-FA-FMK, a potent irreversible inhibitor of cysteine proteases including caspase 3, was also identified to inhibit 3CL^pro^ (IC_50_ = 11.39 μM). The predicted binding models of these inhibitors to 3CL^pro^ showed that they bound to the active site of 3CL^pro^ in the same manner as observed in other peptide-like 3CL^pro^ inhibitors ^2^, suggesting that they share the same mode of action for inhibition of 3CL^pro^ activity.

LLL-12 inhibited the 3CL^pro^ (IC_50_ = 9.84 μM) and it was originally generated by structure-based design targeting the signal transducer and activator of transcription 3 (STAT3) for cancer therapy, where it inhibited STAT3 phosphorylation and induced apoptosis ^27^. Antiviral activity of LLL-12 has been reported against HIV ^28^. The mechanism of action for its antiviral effect was unclear, though it suppressed HIV-1 infection in macrophages ^28^. In our current study, we found that LLL-12 inhibited the SARS-CoV-2 3CL^pro^ activity.

In conclusion, this study employed an enzymatic assay for qHTS that identified 23 SARS-CoV-2 3CL^pro^ inhibitors from a collection of approved drugs, drug candidates, and bioactive compounds. These 3CL^pro^ inhibitors can be combined with drugs of different targets to evaluate their potential in drug cocktails for the treatment of COVID-19. In addition, they can also serve as starting points for medicinal chemistry optimization to improve potency and drug like properties.

## Materials and Methods

### Materials

3CL^pro^ of SARS-CoV-2 with an N-terminal MBP-tag, sensitive internally quenched fluorogenic substrate, and assay buffer were obtained from BPS Bioscience (San Diego, CA, USA). The enzyme was expressed in E. coli expression system, has a molecular weight of 77.5 kDa. The peptide substrate contains 14 amino sequence (KTSAVLQSGFRKME) with Dabcyl and Edans attached on its N- and C-terminals, respectively. The reaction buffer is composed of 20 mM Tris-HCl (pH 7.3), 100 mM NaCl, 1 mM EDTA, 0.01 % BSA (bovine serum albumin), and 1 mM 1,4-dithio-D, L-threitol (DTT). GC376 (CAS No: 1416992-39-6) was purchased from Aobious (Gloucester, MA, USA). Library of Pharmacologically Active Compounds (LOPAC) was purchased from Sigma-Aldrich (St. Louis, MO, USA). All other compound libraries were sourced by the National Center for Advancing Translational Sciences (NCATS) including the NCATS Pharmaceutical Collection (NPC) ^29^, anti-infective, MIPE5.0, and NPACT libraries. LOPAC library has 1280 compounds consisting of marketed drugs and pharmaceutically relevant structures with biological activities. NPC library contains 2552 FDA approved drugs, investigational drugs, animal drugs and anti-infectives. Anti-infective library is a NCATS collection that contains 739 compounds that specifically target to viruses. MIPE 5.0 library includes 2480 compounds that are mixed with approved and investigational compounds, and mechanistic based compounds focusing on oncology. NPACT library contains 5099 structurally diverse compounds consisting of approved drugs, investigational drugs, and natural products.

### 3CL^pro^ enzyme assay

The 3CL^pro^ enzyme assay was developed in 384-well black, medium binding microplates (Greiner Bio-One, Monroe, NC, USA) with a total volume of 20 µL and then miniaturized to 1536-well format. In 384-well plate format, 10 µL enzyme in reaction buffer was added into each well, followed by the addition of 10 µL substrate. Fluorescent intensity was measured at different time points on a PHERAstar FSX plate reader (BMG Labtech, Cary, NC, USA) with Ex=340 nm/Em=460 nm after the addition of substrate. The experiment was conducted at both room temperature (RT) and 37 °C.

Steady-state kinetic parameters were evaluated using 50 nM 3CL^pro^ and different concentrations of substrate. In brief, 10 µL/well enzyme was added into 384-well plate. The reaction was then initialized by adding the substrate solutions at different concentrations. The substrate stock solution was serially diluted 1:2 to obtain seven concentrations. The final concentrations used in this test were 160, 80, 40, 20, 10, 5, and 2.5 µM. The fluorescent intensity was measured at 5, 10, 15, and 30 min.

### Compound library screening, confirmation and counter screen

For the primary screen, library compounds were formatted in 1536-well plates with 4 compounds concentrations at inter-plate titration of 1:5 with the highest concentration of 10 mM for most of compounds in the libraries. The SARS-CoV-2 3CL^pro^ assay was initiated by dispensing 2 µL/well of 50 nM enzyme solution into 1536 black bottom microplates (Greiner Bio-One, Monroe, NC, USA) by a Multidrop Combi disperser (Thermo Fisher Scientific, Waltham, MA, USA), followed by pin transfer of 23 nL compounds in DMSO solution using an automated pintool workstation (WAKO Scientific Solutions, San Diego, CA). After 30 min incubation at RT, 2 µL/well 20 μM substrate solution was dispensed into the assay plates to initiate the enzyme reaction. After 3 h incubation at RT, the plates were read at 460 nm emission upon excitation at 340 nm.

Following primary screening, selected hit compounds were diluted with intraplate 11-point dilution at 1:3 ratio and tested using the same enzyme assay as the primary screen. Each compound was tested in three biological replicates.

A counter-screen assay to eliminate the fluorescence quenching compounds was carried out by dispensing 4 μL of substrate containing fluorescent Edans fragment, SGFRKME-Edans, into 1536-well assay plates. Compounds were pin transferred as 23 nL/well and the fluorescence signal was read.

### SARS-CoV-2 CPE assay

SARS-CoV-2 CPE assay was conducted at Southern Research Institute (Birmingham, AL) as described in previous reports ^30, 31^. In brief, high ACE2 expressing Vero E6 cells were inoculated with SARS-CoV-2 (USA_WA1/2020) at 0.002 M.O.I. After infection of 72 h at 37 °C and 5% CO_2_, the cell viability was examined with CellTiter-Glo ATP content assay kit (Promega, Madison, WI, USA). CPE raw data were normalized to non-infected cells and virus infected cells only which were set as 100% efficacy and 0 efficacy, respectively. In addition, the compound cytotoxicity was evaluated in the same cells by measuring ATP content in the absence of virus. Compound cytotoxicity raw data were normalized with wells containing cells only as 100 % viability (0% cytotoxicity), and wells containing media only as 0% viability (100% cytotoxicity).

### Modeling

Modeling and docking studies were performed using the Molecular Operating Environment (MOE) program (Chemical Computing Group ULC, Montreal, QC, Canada). The crystal structure of 3CL^pro^ in complex with a peptide-like inhibitor N3 (PDB code 6LU7) ^2^ was used to dock inhibitors to the active site of 3CL^pro^. The ligand induced fit docking protocol was used and the binding affinity was evaluated using the GBVI/WSA score. Covalent docking was performed for inhibitors Z-DEVD-FMK and Z-FA-FMK, with a covalent binding to residue Cyc145. Finally, energy minimization was performed to refine the predicted binding complex.

### Data analysis and statistics

The primary screen data was analyzed using a customized software developed in house at NCATS ^29^. Raw data were normalized to relative controls, in which the DMSO alone was set as 0% inhibitory activity, the reaction buffer containing substrate only was set as 100% inhibitory activity. Concentration-response curves were fitted and IC_50_ values of confirmed compounds were calculated using the GraphPad Prism software (San Diego, CA, USA). Data are presented as mean ± standard deviation (SD). Statistical significance was analyzed using one-way ANOVA and difference was defined as P<0.05.

## ACKNOWLEDGEMENTS

This work was supported by the Intramural Research Programs of the National Center for Advancing Translational Sciences, National Institutes of Health. The authors received no other financial supports for the research, authorship, and/or publication of this article.

## Supplementary Information

**Supplementary Figure 1.**
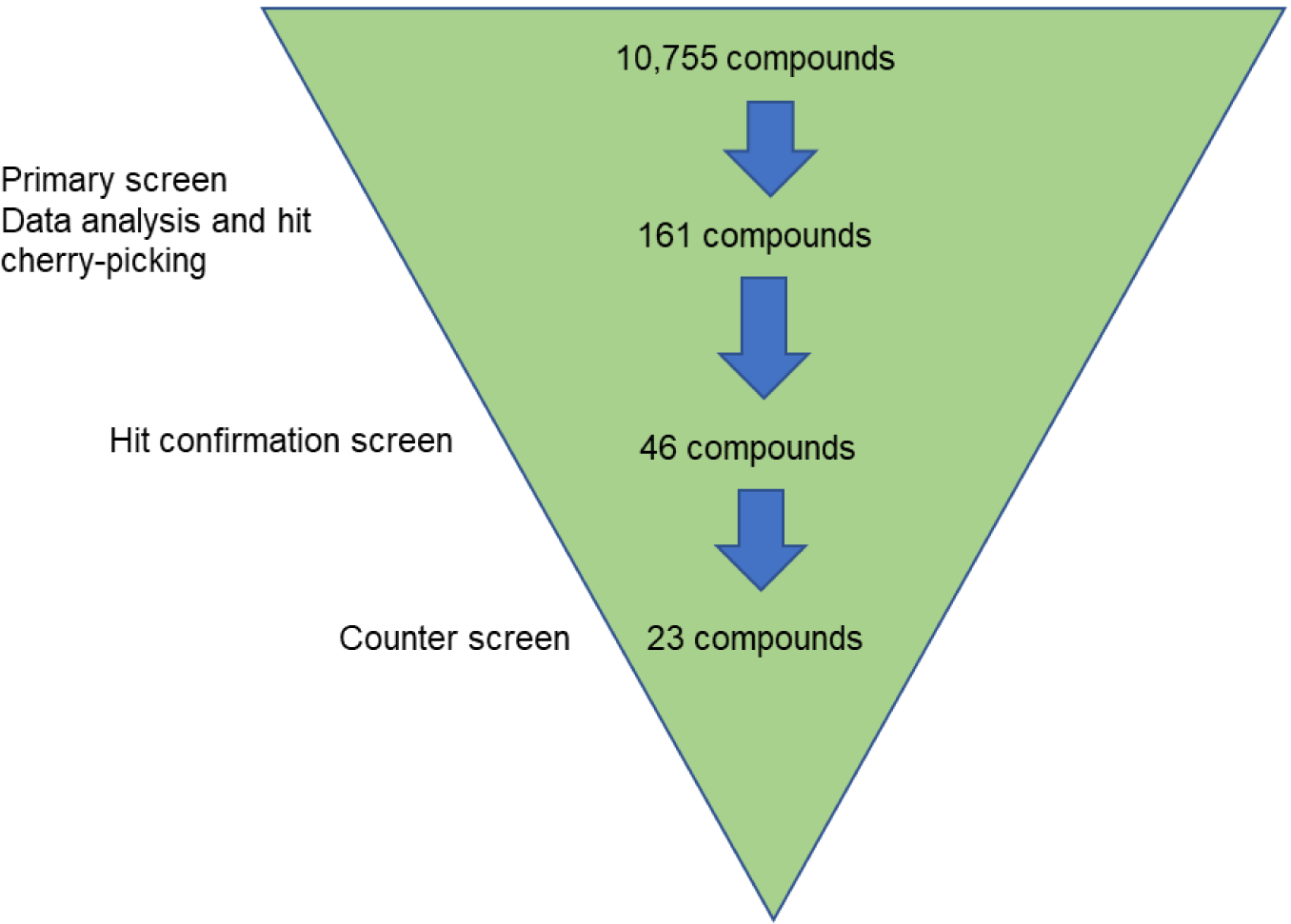
Triage strategy used to narrow down the starting 10,755 small molecules from the primary HTS campaign which led to the identification of 23 compounds having IC50s < 30 μM, maximal inhibition > 60% against SARS-CoV-2 3CL^pro^.

**Supplementary Figure 2.**
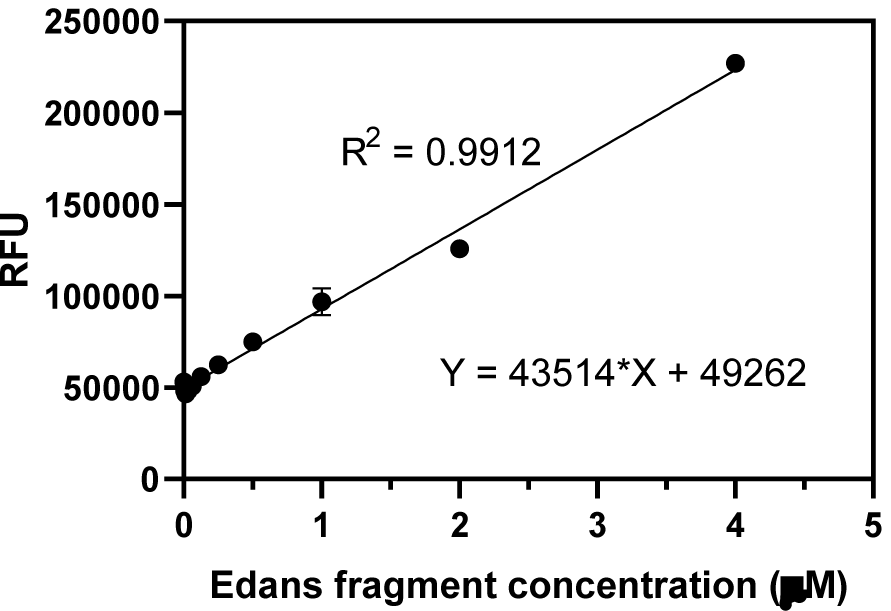
Standard curve generated by diluting SGFRKME-Edans (fluorescent fragment in assay byproducts) in 20 µM substrate solution. The relative fluorescent unit (RFU) is linearly proportional to the amount of SGFRKME-Edans.

**Supplementary Figure 3.**
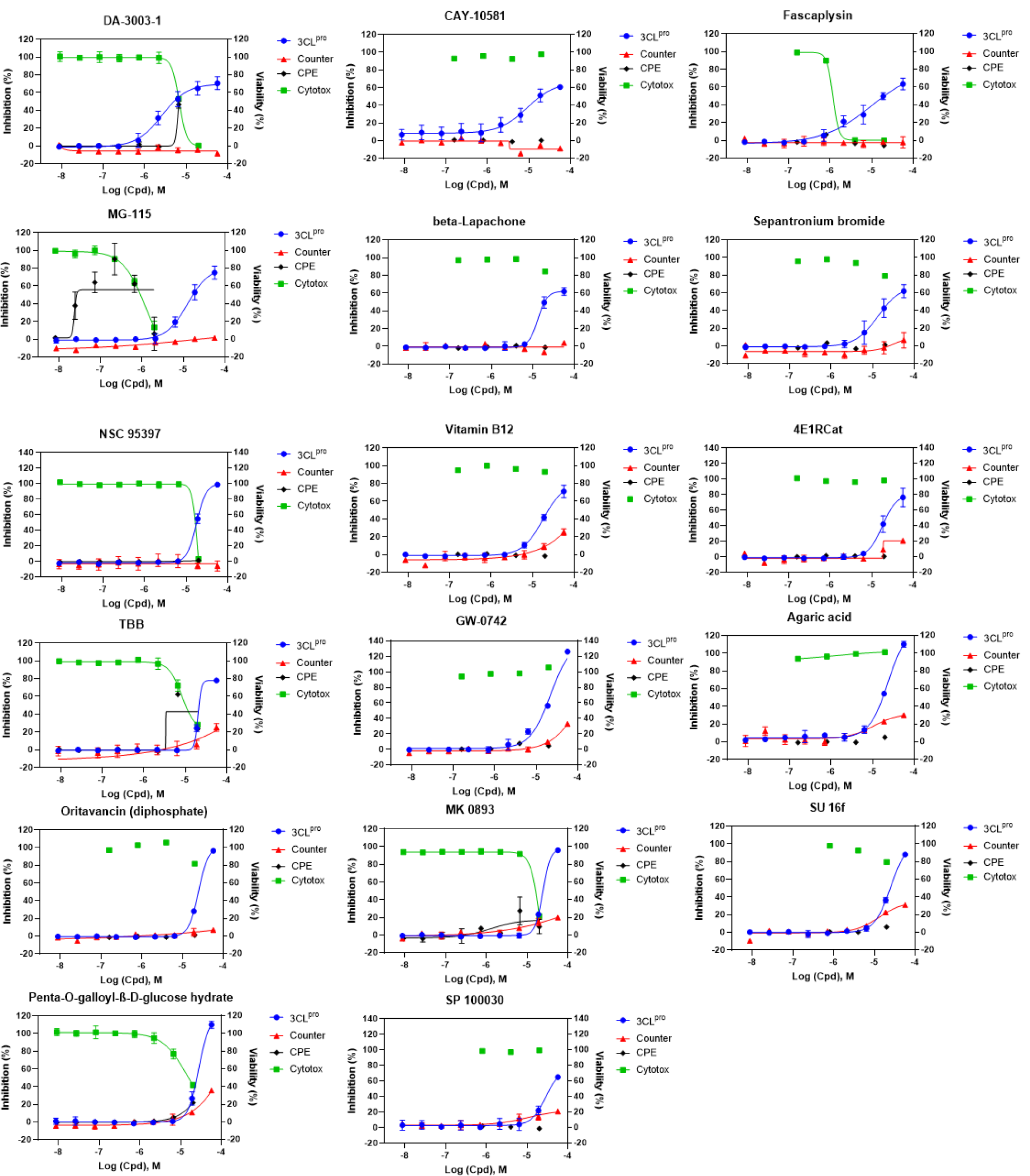
Concentration-response curves of identified compounds by 3CL^pro^ assay. These compounds showed relatively negligible quenching effect relative to enzyme inhibitory activity.

**Supplementary Figure 4.**
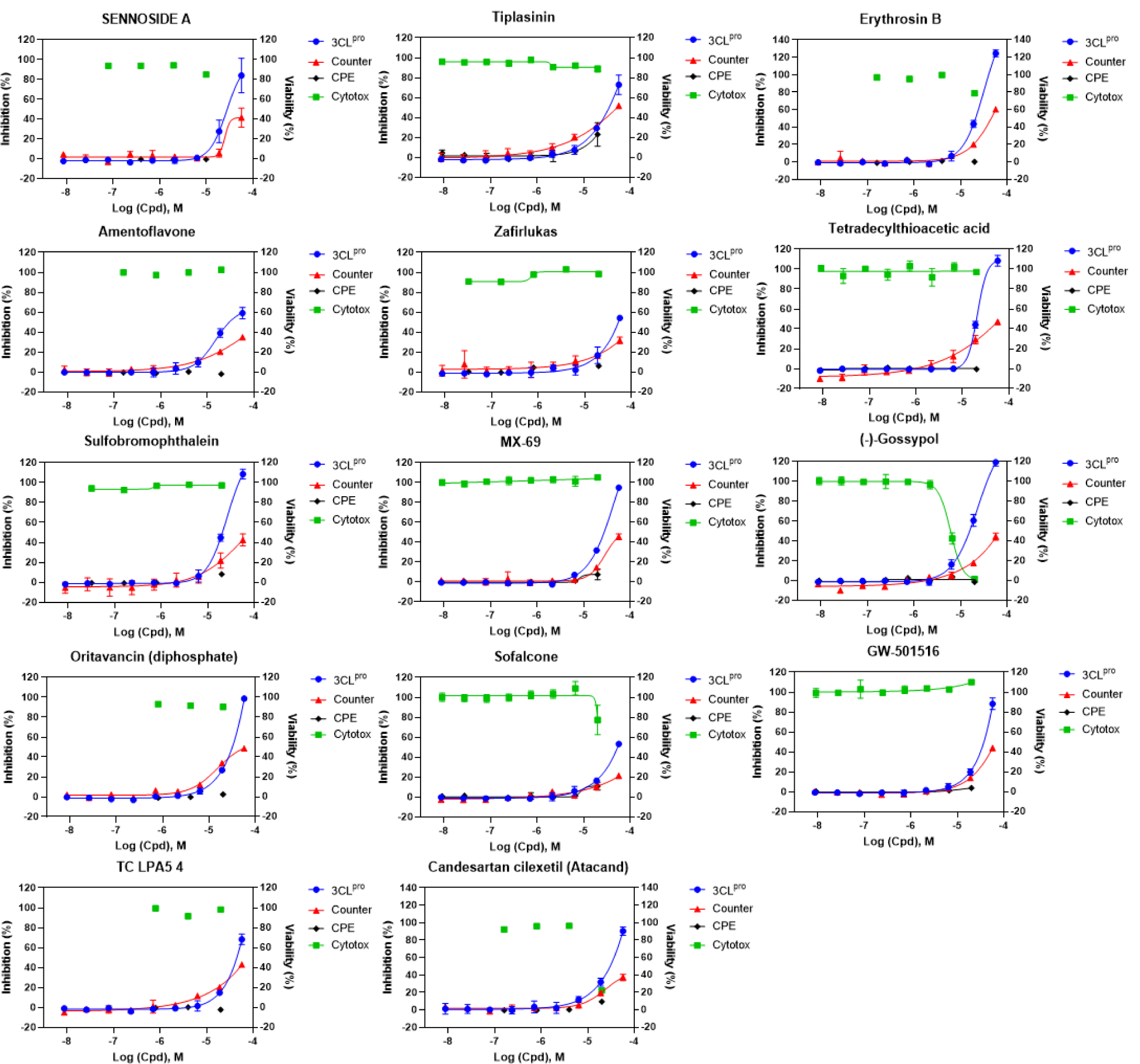
Concentration-response curves of compounds showed apparent quenching effect in counter screen.

## References

1. Wu, F.; Zhao, S.; Yu, B.; Chen, Y.-M.; Wang, W.; Song, Z.-G.; Hu, Y.; Tao, Z.-W.; Tian, J.-H.; Pei, Y.-Y.; Yuan, M.-L.; Zhang, Y.-L.; Dai, F.-H.; Liu, Y.; Wang, Q.-M.; Zheng, J.-J.; Xu, L.; Holmes, E. C.; Zhang, Y.-Z., A new coronavirus associated with human respiratory disease in China. Nature 2020, 579 (7798), 265–269.

2. Jin, Z.; Du, X.; Xu, Y.; Deng, Y.; Liu, M.; Zhao, Y.; Zhang, B.; Li, X.; Zhang, L.; Peng, C.; Duan, Y.; Yu, J.; Wang, L.; Yang, K.; Liu, F.; Jiang, R.; Yang, X.; You, T.; Liu, X.; Yang, X.; Bai, F.; Liu, H.; Liu, X.; Guddat, L. W.; Xu, W.; Xiao, G.; Qin, C.; Shi, Z.; Jiang, H.; Rao, Z.; Yang, H., Structure of Mpro from SARS-CoV-2 and discovery of its inhibitors. Nature 2020, 582 (7811), 289–293.

3. Thiel, V.; Ivanov, K. A.; Putics, Á.; Hertzig, T.; Schelle, B.; Bayer, S.; Weißbrich, B.; Snijder, E. J.; Rabenau, H.; Doerr, H. W.; Gorbalenya, A. E.; Ziebuhr, J., Mechanisms and enzymes involved in SARS coronavirus genome expression. Journal of General Virology 2003, 84 (9), 2305–2315.

4. Zhang, L.; Lin, D.; Sun, X.; Curth, U.; Drosten, C.; Sauerhering, L.; Becker, S.; Rox, K.; Hilgenfeld, R., Crystal structure of SARS-CoV-2 main protease provides a basis for design of improved α-ketoamide inhibitors. Science 2020, 368 (6489), 409.

5. Anand, K.; Ziebuhr J Fau - Wadhwani, P.; Wadhwani P Fau - Mesters, J. R.; Mesters Jr Fau - Hilgenfeld, R.; Hilgenfeld, R., Coronavirus main proteinase (3CLpro) structure: basis for design of anti-SARS drugs. Science 2003, 300 (5626), 1763–1767.

6. De Clercq, E.; Li, G., Approved Antiviral Drugs over the Past 50 Years. Clin Microbiol Rev 2016, 29 (3), 695–747.

7. Blanchard, J. E.; Elowe, N. H.; Huitema, C.; Fortin, P. D.; Cechetto, J. D.; Eltis, L. D.; Brown,E. D., High-throughput screening identifies inhibitors of the SARS coronavirus main proteinase. Chem Biol 2004, 11 (10), 1445–1453.

8. Inglese, J.; Auld, D. S.; Jadhav, A.; Johnson, R. L.; Simeonov, A.; Yasgar, A.; Zheng, W.; Austin, C. P., Quantitative high-throughput screening: A titration-based approach that efficiently identifies biological activities in large chemical libraries. Proceedings of the National Academy of Sciences 2006, 103 (31), 11473.

9. Copeland, R. A., Mechanistic considerations in high-throughput screening. Anal Biochem 2003, 320 (1), 1–12.

10. Ma, C.; Sacco, M. D.; Hurst, B.; Townsend, J. A.; Hu, Y.; Szeto, T.; Zhang, X.; Tarbet, B.; Marty, M. T.; Chen, Y.; Wang, J., Boceprevir, GC-376, and calpain inhibitors II, XII inhibit SARS-CoV-2 viral replication by targeting the viral main protease. Cell Research 2020.

11. Wang, Y.; Jadhav, A.; Southal, N.; Huang, R.; Nguyen, D.-T., A grid algorithm for high throughput fitting of dose-response curve data. Curr Chem Genomics 2010, 4, 57–66.

12. Brimacombe, K. R.; Zhao, T.; Eastman, R. T.; Hu, X.; Wang, K.; Backus, M.; Baljinnyam, B.; Chen, C. Z.; Chen, L.; Eicher, T.; Ferrer, M.; Fu, Y.; Gorshkov, K.; Guo, H.; Hanson, Q. M.; Itkin, Z.; Kales, S. C.; Klumpp-Thomas, C.; Lee, E. M.; Michael, S.; Mierzwa, T.; Patt, A.; Pradhan, M.; Renn, A.; Shinn, P.; Shrimp, J. H.; Viraktamath, A.; Wilson, K. M.; Xu, M.; Zakharov, A. V.; Zhu, W.; Zheng, W.; Simeonov, A.; Mathé, E. A.; Lo, D. C.; Hall, M. D.; Shen, M., An OpenData portal to share COVID-19 drug repurposing data in real time. bioRxiv 2020, 2020.06.04.135046.

13. Zumla, A.; Chan, J. F. W.; Azhar, E. I.; Hui, D. S. C.; Yuen, K.-Y., Coronaviruses - drug discovery and therapeutic options. Nature reviews. Drug discovery 2016, 15 (5), 327–347.

14. Cao, B.; Wang, Y.; Wen, D.; Liu, W.; Wang, J.; Fan, G.; Ruan, L.; Song, B.; Cai, Y.; Wei, M.; Li, X.; Xia, J.; Chen, N.; Xiang, J.; Yu, T.; Bai, T.; Xie, X.; Zhang, L.; Li, C.; Yuan, Y.; Chen, H.; Li, H.; Huang, H.; Tu, S.; Gong, F.; Liu, Y.; Wei, Y.; Dong, C.; Zhou, F.; Gu, X.; Xu, J.; Liu, Z.; Zhang, Y.; Li, H.; Shang, L.; Wang, K.; Li, K.; Zhou, X.; Dong, X.; Qu, Z.; Lu, S.; Hu, X.; Ruan, S.; Luo, S.; Wu, J.; Peng, L.; Cheng, F.; Pan, L.; Zou, J.; Jia, C.; Wang, J.; Liu, X.; Wang, S.; Wu, X.; Ge, Q.; He, J.; Zhan, H.; Qiu, F.; Guo, L.; Huang, C.; Jaki, T.; Hayden, F. G.; Horby, P. W.; Zhang, D.; Wang, C., A Trial of Lopinavir-Ritonavir in Adults Hospitalized with Severe Covid-19. N Engl J Med 2020, 382 (19), 1787–1799.

15. Dai, W.; Zhang, B.; Jiang, X.-M.; Su, H.; Li, J.; Zhao, Y.; Xie, X.; Jin, Z.; Peng, J.; Liu, F.; Li, C.; Li, Y.; Bai, F.; Wang, H.; Cheng, X.; Cen, X.; Hu, S.; Yang, X.; Wang, J.; Liu, X.; Xiao, G.; Jiang, H.; Rao, Z.; Zhang, L.-K.; Xu, Y.; Yang, H.; Liu, H., Structure-based design of antiviral drug candidates targeting the SARS-CoV-2 main protease. Science 2020, 368 (6497), 1331.

16. Gotoh, Y.; Doi, A.; Furuta, E.; Dubrac, S.; Ishizaki, Y.; Okada, M.; Igarashi, M.; Misawa, N.; Yoshikawa, H.; Okajima, T.; Msadek, T.; Utsumi, R., Novel antibacterial compounds specifically targeting the essential WalR response regulator. The Journal of Antibiotics 2010, 63 (3), 127–134.

17. Weinberg, J. B.; Shugars, D. C.; Sherman, P. A.; Sauls, D. L.; Fyfe, J. A., Cobalamin Inhibition of HIV-1 Integrase and Integration of HIV-1 DNA into Cellular DNA. Biochemical and Biophysical Research Communications 1998, 246 (2), 393–397.

18. Li, D.; Lott, W. B.; Martyn, J.; Haqshenas, G.; Gowans, E. J., Differential Effects on the Hepatitis C Virus (HCV) Internal Ribosome Entry Site by Vitamin B_12_ and the HCV Core Protein. Journal of Virology 2004, 78 (21), 12075.

19. Small, E. J.; Halabi, S.; Ratain, M. J.; Rosner, G.; Stadler, W.; Palchak, D.; Marshall, E.; Rago, R.; Hars, V.; Wilding, G.; Petrylak, D.; Vogelzang, N. J., Randomized Study of Three Different Doses of Suramin Administered With a Fixed Dosing Schedule in Patients With Advanced Prostate Cancer: Results of Intergroup 0159, Cancer and Leukemia Group B 9480. Journal of Clinical Oncology 2002, 20 (16), 3369–3375.

20. Wiedemar, N.; Hauser, D. A.; Mäser, P., 100 Years of Suramin. Antimicrobial Agents and Chemotherapy 2020, 64 (3), e01168–19.

21. Henß, L.; Beck, S.; Weidner, T.; Biedenkopf, N.; Sliva, K.; Weber, C.; Becker, S.; Schnierle, B. S., Suramin is a potent inhibitor of Chikungunya and Ebola virus cell entry. Virol J 2016, 13 (1), 149–149.

22. Mastrangelo, E.; Pezzullo, M.; Tarantino, D.; Petazzi, R.; Germani, F.; Kramer, D.; Robel, I.; Rohayem, J.; Bolognesi, M.; Milani, M., Structure-Based Inhibition of Norovirus RNA-Dependent RNA Polymerases. Journal of Molecular Biology 2012, 419 (3), 198–210.

23. Albulescu, I. C.; van Hoolwerff, M.; Wolters, L. A.; Bottaro, E.; Nastruzzi, C.; Yang, S. C.; Tsay, S.-C.; Hwu, J. R.; Snijder, E. J.; van Hemert, M. J., Suramin inhibits chikungunya virus replication through multiple mechanisms. Antiviral Research 2015, 121, 39–46.

24. da Silva, C. S. B.; Thaler, M.; Tas, A.; Ogando, N. S.; Bredenbeek, P. J.; Ninaber, D. K.; Wang, Y.; Hiemstra, P. S.; Snijder, E. J.; van Hemert, M. J., Suramin inhibits SARS-CoV-2 infection in cell culture by interfering with early steps of the replication cycle. Antimicrobial Agents and Chemotherapy 2020, AAC.00900-20.

25. Knoblach, S. M.; Alroy, D. A.; Nikolaeva, M.; Cernak, I.; Stoica, B. A.; Faden, A. I., Caspase Inhibitor z-DEVD-fmk Attenuates Calpain and Necrotic Cell Death in Vitro and after Traumatic Brain Injury. Journal of Cerebral Blood Flow & Metabolism 2004, 24 (10), 1119–1132.

26. Barut, ᅞ.; Ünlü, Y. A.; Karaoğlan, A.; Tunçdemir, M.; Dağistanli, F. K.; Öztürk, M.; Çolak, A., The neuroprotective effects of z-DEVD.fmk, a caspase-3 inhibitor, on traumatic spinal cord injury in rats. Surgical Neurology 2005, 64 (3), 213–220.

27. Lin, L.; Hutzen, B.; Li, P.-K.; Ball, S.; Zuo, M.; DeAngelis, S.; Foust, E.; Sobo, M.; Friedman, L.; Bhasin, D.; Cen, L.; Li, C.; Lin, J., A novel small molecule, LLL12, inhibits STAT3 phosphorylation and activities and exhibits potent growth-suppressive activity in human cancer cells. Neoplasia 2010, 12 (1), 39–50.

28. Appelberg, K. S.; Wallet, M. A.; Taylor, J. P.; Cash, M. N.; Sleasman, J. W.; Goodenow, M. M., HIV-1 Infection Primes Macrophages Through STAT Signaling to Promote Enhanced Inflammation and Viral Replication. AIDS Research and Human Retroviruses 2017, 33 (7), 690–702.

29. Huang, R.; Zhu, H.; Shinn, P.; Ngan, D.; Ye, L.; Thakur, A.; Grewal, G.; Zhao, T.; Southall, N.; Hall, M. D.; Simeonov, A.; Austin, C. P., The NCATS Pharmaceutical Collection: a 10-year update. Drug Discovery Today 2019, 24 (12), 2341–2349.

30. Gorshkov, K.; Chen, C. Z.; Bostwick, R.; Rasmussen, L.; Xu, M.; Pradhan, M.; Tran, B. N.; Zhu, W.; Shamim, K.; Huang, W.; Hu, X.; Shen, M.; Klumpp-Thomas, C.; Itkin, Z.; Shinn, P.; Simeonov, A.; Michael, S.; Hall, M. D.; Lo, D. C.; Zheng, W., The SARS-CoV-2 cytopathic effect is blocked with autophagy modulators. bioRxiv 2020, 2020.05.16.091520.

31. Chen, C. Z.; Xu, M.; Pradhan, M.; Gorshkov, K.; Petersen, J.; Straus, M. R.; Zhu, W.; Shinn, P.; Guo, H.; Shen, M.; Klumpp-Thomas, C.; Michael, S. G.; Zimmerberg, J.; Zheng, W.; Whittaker, G. R., Identifying SARS-CoV-2 entry inhibitors through drug repurposing screens of SARS- S and MERS-S pseudotyped particles. bioRxiv 2020, 2020.07.10.197988.

